# Altered hippocampal activation in seizure-prone *CACNA2D2* knockout mice

**DOI:** 10.1101/2023.11.08.565511

**Authors:** Alyssa Danis, Ashlynn A. Gallagher, Ashley N. Anderson, Arielle Isakharov, Kathleen A. Beeson, Eric Schnell

**Affiliations:** Department of Anesthesiology and Perioperative Medicine, Oregon Health & Science University, Portland, OR, 97239; Research and Development Service, Portland VA Health Care System, Portland, OR, 97239, Portland, OR, 97239; Neuroscience Graduate Program, Oregon Health & Science University, Portland, OR, 97239

## Abstract

The voltage-gated calcium channel subunit α2δ-2 controls calcium-dependent signaling in neurons, and loss of this subunit causes epilepsy in both mice and humans. To determine whether mice without α2δ-2 demonstrate hippocampal activation or histopathological changes associated with seizure activity, we measured expression of the activity-dependent gene *c-fos* and various histopathological correlates of temporal lobe epilepsy in hippocampal tissue from wildtype (WT) and α2δ-2 knockout (*CACNA2D2* KO) mice using immunohistochemical staining and confocal microscopy. Both genotypes demonstrated similarly sparse *c-fos* expression within the hippocampal dentate granule cell layer (GCL) at baseline, consistent with no difference in basal activity of granule cells between genotypes. Surprisingly, when mice were assayed 1 hour after handling-associated convulsions, KO mice had fewer c-fos-positive cells in the dentate gyrus, indicating that activity in the dentate gyrus actually decreased. However, the dentate was significantly more active in KO mice compared to WT after administration of a subthreshold pentylenetetrazole dose, consistent with increased susceptibility to proconvulsant stimuli. Other histopathological markers of temporal lobe epilepsy in these mice, including markers of neurogenesis, glial activation, and mossy fiber sprouting, were similar in WT and KO mice, apart from a small but significant increase in hilar mossy cell density, opposite to what is typically found in mice with temporal lobe epilepsy. This suggests that the differences in seizure-associated hippocampal function in the absence of α2δ-2 protein are likely due to altered functional properties of the network without associated structural changes in the hippocampus at the typical age of seizure onset.

**Significance Statement:** Calcium channel α2δ subunits play important roles in controlling neuronal circuit structure and function, and mutation of the α2δ-2 isoform of this protein is associated with spontaneous seizures. In this study, we find that seizures in α2δ-2 mutant mice involve altered hippocampal activation without substantial histopathological changes in hippocampal structure. This suggests that the differences in seizure-associated hippocampal function are likely due to altered functional properties of the network as well as to the contribution of additional brain regions to seizures in mice lacking this protein.

## Introduction

Mutations in ion channel genes can dramatically alter neuronal function, and in severe cases cause genetic forms of epilepsy known as channelopathies (Davies & Hanna, 1999). In particular, voltage-gated calcium channels (VGCCs) play critical roles in controlling neuronal excitability, synaptic transmission, and synaptic plasticity, and mutations in many VGCC genes cause treatment-resistant forms of epilepsy that can begin neonatally (Catterall, 2011). These syndromes are sometimes associated with structural brain defects, which could alter neuronal circuit connectivity and thus secondarily contribute to circuit hyperexcitability (Rajakulendran & Hanna, 2016).

Although channelopathies often result from mutations in pore-forming subunits that critically alter channel properties, mutations in non-pore-forming auxiliary VGCC subunits also cause genetic forms of epilepsy. One such subunit is the α2δ-2 protein, encoded by the *CACNA2D2* gene, originally associated with a heritable form of epilepsy in the spontaneous α2δ-2 mutant *ducky* mouse (Barclay et al., 2001; Snell, 1955). Subsequent work has identified additional spontaneous mutant *CACNA2D2* alleles in mice (Brill et al., 2004), and epileptic phenotypes have been associated with both targeted mutations of *CACNA2D2* and spontaneous *CACNA2D2* mutations in humans (Edvardson et al., 2013; Ivanov et al., 2004; Pippucci et al., 2013).

Although the α2δ-2 subunit does not notably alter VGCC voltage-dependence or conductance, it does play a significant role in the surface and subcellular trafficking of VGCCs, which could dramatically alter neuronal circuit structure and function (Barclay et al., 2001). Additionally, a separate member of the α2δ family of proteins, the α2δ-1 isoform, is a neuronal thrombospondin receptor with synaptogenic functions, demonstrating that α2δ subunits can have signaling roles independent of VGCCs that could alter neuronal circuits (Eroglu et al., 2009). However, despite the known and hypothesized cellular roles of α2δ-2, the specific circuit alterations in *CACNA2D2* mutant mice that contribute to seizure initiation and propagation are unknown.

*CACNA2D2* mutant mice exhibit handling-associated convulsions beginning at an early age, and electroencephalographic spike-wave discharge (SWD) activity and absence-type seizures in young adult mice, which can progress to generalized seizures that are strongly associated with high juvenile mortality (Barclay et al., 2001; Meier, 1968; Snell, 1955). SWDs are typically associated with aberrant thalamocortical activation (Blumenfeld, 2005), but as α2δ-2 protein is expressed in the hippocampus (Cole et al., 2005; Lein et al., 2007), we wondered whether structural or functional alterations in the hippocampus might be associated with, and potentially contribute to, the progression to generalized seizure activity in *CACNA2D2* mutant mice. To investigate a potential relationship between *CACNA2D2*-associated epilepsy and structural brain changes associated with temporal lobe epilepsy (TLE), we examined hippocampal sections from WT and *CACNA2D2* KO mice for patterns of granule cell activity based on expression of the activity-dependent immediate early gene *c-fos*, as well as for a range of other histopathological markers typically associated with TLE.

## Methods

### Mice

*CACNA2D2* KO mice (*CACNA2D2^tm1Svi^*, MGI:3055290, RRID:MGI:3055322) were created by homologous recombination and maintained on a C57BL/6 background (Ivanov et al., 2004). Mice were housed in an AAALAC-certified facility, and all animal procedures were performed in accordance with the Portland VA Health Care System’s animal care committee’s regulations. Mice were placed on a 12 hour light-dark cycle with continuous access to food/water *ad libitum*, and experiments/harvests were performed at Zeitgeber time ZT4-6 (4-6 hours after lights on). This strain was maintained by breeding heterozygotic mice, which produced WT and KO littermates for experimental analysis. All mice were genotyped by PCR as previously described (Beeson et al., 2020), and all KOs were additionally verified by morphological phenotype, which consisted of small size, hind-limb clasping, and ataxic gait. Mice were group housed with same-sex littermates of mixed genotype at weaning.

WT and KO mice of both sexes were analyzed at 4-5 weeks of age. In a majority of cases, each mouse was paired with a littermate of the converse genotype, but for some litters a Mendelian ratio was not achieved, so mice were paired across litters when necessary to balance genotypes. Given the high mortality of mice at weaning age (∼28do) with conventional husbandry (Meier, 1968), we adapted our weaning protocols to provide soft food continuously after weaning, with gentle handling, as previously shown to improve survival (Geisler et al., 2021). Data from male and female mice in each group were compared to assess for possible sex differences, which were not detected in any of our assays, and thus data from both sexes were combined.

### Acute seizure modeling

Pentylenetetrazole (PTZ; Sigma) was diluted in sterile water at 3 mg/ml, and administered to a subset of mice by intraperitoneal injection at a dosage of 30 mg/kg. Mice were continually observed for seizures for one hour after administration, which were scored on a modified Racine scale for mice (Shibley & Smith, 2002), with a zero score given for normal behavior without any tremor or freezing, and a six for high amplitude jumping/popcorn behavior. After one hour, mice were humanely euthanized by anesthetic overdose for brain harvest.

To assess hippocampal involvement in handling-induced seizure activity (Meier, 1968; Snell, 1955), a separate cohort of WT and KO mice were manually removed from their cages by a gloved investigator by the base of the tail, rotated 180 degrees, and placed on a wire cage top. They were then picked up using a gentle scruffing technique and held in the supine (belly up) position for 5 seconds. After this maneuver, mice were returned to their home cages for 1 hour of observation and seizure scoring prior to euthanasia for histologic analysis.

### Histology

Prior to brain harvest, mice were euthanized under deep isoflurane/avertin anesthesia by transcardiac perfusion and decapitation in accordance with IACUC-approved protocols. Transcardiac perfusion consisted of 5 ml phosphate-buffered saline (PBS; pH 7.4), followed by 10 ml 4% paraformaldehyde in PBS (PFA). After dissection, brains were post-fixed overnight in PFA, rinsed x3 in PBS, and cut into 100 µm thick coronal sections on a Leica VT1000 vibratome.

Free floating brain sections were permeabilized and blocked in 0.4% triton X-100 in PBS (PBST) containing 10% goat serum for 1 hour, and stained overnight at 4C with the following antibodies diluted in PBST + 1.5% goat serum: rabbit anti-c-fos, CellSignaling 2250, RRID:AB_2247211, 1:300; guinea pig anti-ZnT3, Synaptic Systems 197004, RRID:AB_2189667, 1:200; mouse anti-PV, Neuromab 75-455, RRID:AB_2629420, 1:20; goat anti-calretinin, Swant CG1, RRID:AB_10000342, 1:500; rabbit anti-Ki67, Millipore AB9260, RRID:AB_2142366, 1:500; guinea pig anti-Dcx, Millipore AB2253, RRID:AB_1586992, 1:500; mouse anti-GFAP, Sigma G3893, RRID:AB_477010, 1:500; rabbit anti-Iba1, Wako 019-19741, RRID:AB_839504, 1:1000; rat anti-Mac-2, Cedarlane CL8942AP, RRID:AB_10060357, 1:1000. The following morning, slices were rinsed in PBST 3x for 5 minutes each, and incubated in Alexafluor-conjugated secondary antibodies (goat or donkey, Alexa 488/568/647, 1:400 each, Invitrogen) in PBST + 1.5% goat serum at room temperature for 4-6 hrs. Sections were subsequently rinsed, counterstained with DAPI 1:20,000 for 10 minutes, and mounted on glass slides with Fluoromount G. For anti-c-fos, anti-Ki67, anti-Dcx, and anti-GFAP stains, slices underwent an antigen retrieval step at 95C (for 30 minutes followed by gradual cooling per the manufacturer’s instructions; DAKO) prior to permeabilization and blocking to improve antigenicity.

### Imaging and Analysis

Slides were imaged on an upright Scientifica Slicescope with Olympus optics, using 10x 0.4NA, 20x 0.8NA, and 40x 1.4NA objectives, and acquired with a Photometrics 95 camera through a Yokogawa CSU-WI spinning disk confocal system running Slidebook software (Intelligent Imaging, Inc.). After acquisition, images were transferred to a separate computer for manual counting/analysis using FIJI/ImageJ software. WT and KO sections were stained side-by-side with identical solutions for each assay. At mounting, all slides were coded to de-identify animal ID/genotype prior to imaging, and all imaging, image analysis, and quantification was performed by an investigator blinded to animal/slide identity/genotype. Results were unblinded and combined by a 3^rd^ investigator only after analysis and quantification was complete. Cell densities were quantified using either single confocal sections or confocal image Z-stacks, as detailed for each specific experiment below.

c-fos+ cell densities were calculated from confocal image stacks acquired with a 10x objective from up to 4 sections per mouse. For each mouse, c-fos+ cell densities from each image were averaged to obtain a single value per animal. All c-fos+ cells in the GCL or at the GCL/inner molecular layer (IML) border were counted and normalized to the GCL volume in the image stack. GCL volume (in mm^3^) was calculated as the GCL area determined by tracing DAPI-stained granule cell nuclei x depth of stack. To subcategorize putative semilunar granule cells, cells that were outside of the GCL but within 10 µm (approximately one granule cell soma diameter) of the GCL-IML border were counted separately and normalized to the same GCL volume. Data shown in Figure 1 represent the combined cell counts of putative granule cells and semilunar cells in each group; subcategorized data is in the text.

**Figure 1.**
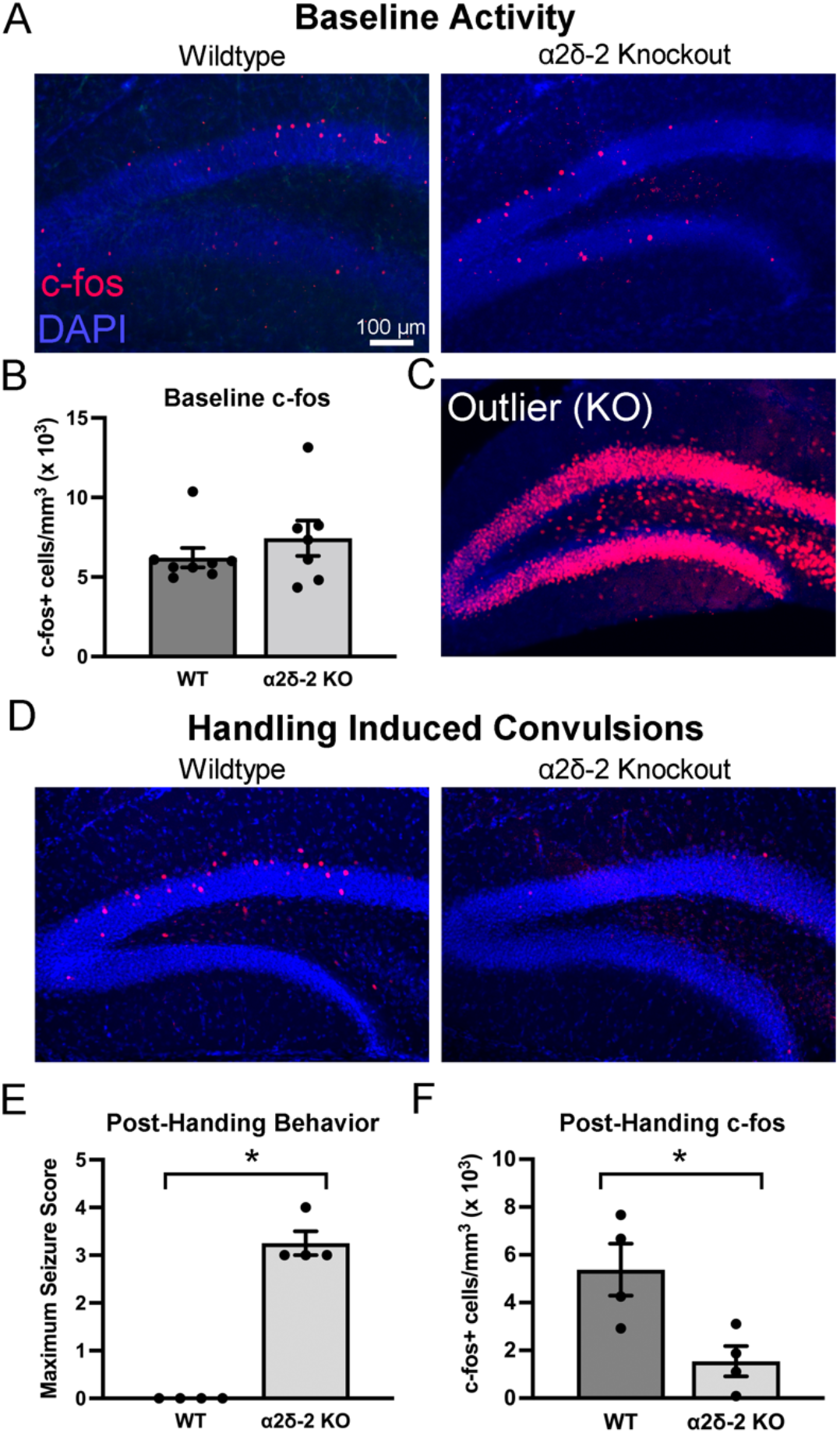
Granule cell expression of the activity-dependent gene *c-fos*. A. Example images from 4 week old WT and *CACNA2D2* KO mouse hippocampal sections, demonstrating c-fos+ granule cells (red) sparsely located throughout the granule cell layer and near the granule cell-molecular layer border. Cell nuclei visualized using DAPI stain (blue). B. Quantification of c-fos+ density (n = 8, 7; p = 0.40). C. One KO mouse excluded from baseline analysis, due to global dentate *c-fos* expression, suggestive of recent seizure. D. Images of *c-fos* expression one hour after gentle handling in representative WT and KO mice. E. Gentle handling caused convulsive seizures in *CACNA2D2* KO, but not WT, mice (n = 4, 4; p < 0.0001). F. Granule cell *c-fos* expression was reduced in *CACNA2D2* mice after handling-induced convulsions (n = 4, 4; p = 0.023).

Hilar mossy cell density was quantified as the number of calretinin-positive cells in the hilar area (bounded by the subgranular zone and edges of the image) divided by the hilar area imaged in 2 dimensions, from a single confocal section acquired at 20x. Parvalbumin-positive (PV+) cell density was quantified as the number of PV+ cells divided by the volume of GCL or CA3 pyramidal cell layer imaged for measurements made in the dentate or CA3 respectively. CA3 pyramidal cell layer width was defined as the average transverse width of the cell layer (defined using DAPI+ nuclei) obtained by measuring at three points along CA3a/b. Mossy fiber sprouting was qualitatively evaluated (blindly) based on the ZnT3 staining pattern, using a modified scoring system (based roughly on the Timm scoring system of (Tauck & Nadler, 1985)) as follows: 0 = no evidence for sprouting, 1 = rare ZnT3+ stain in GCL only, 2 = some ZnT3+ in the GCL with rare ZnT3 in the IML, and 3 = clear ZnT3+ puncta in IML. In no image from either genotype did there appear to be more than 3 putative ZnT3+ terminals in the GCL or IML. Ki67 staining was quantified from confocal stacks obtained using a 40x objective, and all Ki67+ nuclei within 10 µm of the sub-granular zone (SGZ) were included in the analysis, and normalized to GCL volume. Doublecortin (Dcx) staining was very dense in all mice of this age, and quantified by measuring average fluorescence intensity of Dcx+-stained tissue within the GCL, from a single 40x confocal section imaged near the surface of the slice, using identical side-by-side staining and imaging protocols. Astrocytes were quantified from 15 µm tall image stacks obtained of the granule cell and molecular layers using a 40x objective, and defined as GFAP+ processes emanating from a clear central location (soma) that was within the imaged volume. Microglia were quantified from single confocal sections obtained at 40x, as cells with a clear soma present within the image based on Iba1 staining pattern. Due to low density of positive cells, Mac-2+ cells were quantified from confocal image stacks of the dentate, including molecular layer, GCL and hilus, acquired at 20x and normalized to dentate volume imaged.

### Statistics

Images were acquired from up to 4 hippocampal sections per animal for each experiment; sections were excluded (prior to unblinding) if the section was damaged during mounting or an air bubble obscured tissue visualization. The quantified data from each of these sections were averaged to provide one data point for each measurement per animal (n=1). A separate investigator unblinded these aggregate values and averaged them for each genotype, thus n = the number of mice analyzed for each measurement. Data for each group are presented ± Standard Error of the Mean (SEM).

Statistical analyses were performed using Prism software (Graphpad), and statistical information for each comparison are presented in Table 1. For continuous numerical data (cell densities, layer width, fluorescence), normality of each data set was first determined by Shapiro-Wilk test. Group means were compared using two-tailed unpaired t-tests for normally distributed data, and using a Mann-Whitney non-parametric test for non-normally distributed data, with significance set at p < 0.05. Additionally detailed methods, raw data, and processed data will be provided upon request.

**Table 1:** Statistical Table.

| **Comparison** | **Data structure** | **Statistical test** | **Difference between means** | **Power (95% confidence intervals)** |
| --- | --- | --- | --- | --- |
| Baseline c-fos+ total Cell Density | Non-Normal distribution (W = 0.6411) | Mann-Whitney test | 1.603 (medians) | Rank sums 56 vs. 64 |
| Baseline c-fos+ total Cell Density (including outlier) | Non-Normal distribution (W = 0.4324) | Mann-Whitney test | 1.964 (medians) | Rank sums 56 vs. 80 |
| Baseline c-fos+ Semi-lunar cell density | Non-Normal distribution (W = 0.6780) | Mann-Whitney test | 19.14 (medians) | Rank sums 58 vs 62 |
| Baseline c-fos+ Granule Cell Density | Non-Normal distribution (W = 0.6960) | Mann-Whitney test | 1649 (medians) | Rank sums 57 vs 63 |
| Handling Seizure Score | Normal distribution | Unpaired t-test | 3.250 | 2.638 to 3.862 |
| Handling c-fos+ Cell Density (WT vs. KO) | Normal distribution | Unpaired t-test | -3.834 | -6.919 to 0.7493 |
| WT: Baseline vs Handled c-fos+ Cell Density | Non-Normal distribution (W = 0.6411) | Mann-Whitney test | -0.278 (medians) | Rank sums 54 vs. 24 |
| KO: Baseline vs Handled c-fos+ Cell Density | Normal distribution | Unpaired t-test | -5.895 | -9.456 to -2.334 |
| PTZ Seizure Score | Normal distribution | Unpaired t-test | 4.375 | 3.035 to 5.715 |
| PTZ c-fos+ Cell Density | Non-Normal distribution (W = 0.7440) | Mann-Whitney test | 71.40 (medians) | Rank sums 47 vs 89 |
| Calretinin+ Cell Density | Normal distribution | Unpaired t-test | 107.3 | 7.566 to 207.0 |
| CA3 Pyramidal Cell layer thickness | Non-Normal distribution (W = 0.575) | Mann-Whitney test | 5.75 (medians) | Rank sums 38 vs. 40 |
| CA3 Pyramidal Cell Density | Normal distribution | Unpaired t-test | 47.00 | -88.44 to 182.4 |
| Mossy fiber sprouting | Non-Normal distribution (W = 0.7013) | Mann-Whitney test | -.375 (medians) | Rank sums 51.5, 26.5 |
| Ki67+ Cell Density | Normal distribution | Unpaired t-test | 13.61 | -3.455 to 30.67 |
| DCX Intensity | Normal distribution | Unpaired t-test | -120.0 | -585.3 to 345.4 |
| PV+ Cell Density in GCL | Normal distribution | Unpaired t-test | 356.3 | -348.6 to 1061 |
| PV+ Cell Density in hilus | Normal distribution | Unpaired t-test | 4.014 | -196.5 to 204.6 |
| PV+ Cell Density in CA3 | Normal distribution | Unpaired t-test | 3.285 | -4.953 to 11.52 |
| GFAP+ Cell Density | Normal distribution | Unpaired t-test | 2.460 | -2.207 to 7.126 |
| GFAP Intensity | Normal distribution | Unpaired t-test | 8.395 | -38.25 to 55.04 |
| Iba1 Cell Density | Normal distribution | Unpaired t-test | 2.630 | -36.21 to 41.47 |
| Mac-2 Cell Density | Normal distribution | Unpaired t-test | 524 | -178 to 1225 |

## Results

### Juvenile *CACNA2D2* KO mice do not have increased granule cell activity at baseline

*CACNA2D2* mutant mice begin to experience spontaneous generalized convulsions at approximately weaning age (3-4 weeks), which manifest as excited jumping/falling, reeling to one side, stiffening of hindlimbs and tail, and periodic paraplegia (Meier, 1968). To assess whether this phenotype was associated with chronically elevated neuronal activity in the hippocampus, we stained the hippocampal dentate gyrus harvested from 1 month old mice for the activity-dependent immediate early gene *c-fos*. Transcription and translation of *c-fos* is transiently increased after neuronal firing, and thus anti-c-fos staining serves as a reliable histochemical marker of recent neuronal activity (Bullitt, 1990; Morgan et al., 1987). We focused on the dentate gyrus, which serves as the gateway for electrical activity to enter the hippocampus and has very sparse baseline activity in healthy mice (Jung & McNaughton, 1993).

In the dentate, both WT and *CACNA2D2* KO mice had similarly sparse densities of c-fos+ cells within and immediately adjacent to the granule cell layer (Figures 1A-B). Although not quantified explicitly, the density of c-fos+ cells was anecdotally greater in the suprapyramidal blade of the dentate (closer to CA1) than the infrapyramidal blade in both genotypes (as seen in Figure 1A). Many c-fos+ cells were also found along the border of the granule cell layer (GCL) and inner molecular layer (IML), the typical location for semilunar granule cells (Williams et al., 2007). To examine whether these cells might have a selective activation pattern, we separately categorized the c-fos+cells within one cell body diameter of the GCL-IML border as putative semilunar cells, and the c-fos+ cells fully within the GCL as putative granule cells. c-fos+ cell density was not different between genotypes in either location (putative semilunar cells: WT = 862 ± 133 cells/mm^3^, KO = 1185 ± 325 cells/mm^3^, n = 8,7 mice, p = 0.54; putative granule cells: WT = 5350 ± 530 cells/mm^3^, KO = 6250 ± 810 cells/mm^3^, n = 8,7 mice, p = 0.46). These data suggest that at baseline, there is no difference in hippocampal activity in *CACNA2D2* KO mice.

In the above analyses, a single KO mouse was excluded from the group data, as its c-fos+ cell density was over 100 standard deviations above the mean value for the rest of the group, with essentially every single dentate granule cell staining as c-fos+ (Figure 1C). Based on this expression pattern, this mouse almost definitely had a seizure which had propagated through the dentate gyrus in the hour prior to fixation (Barone et al., 1993; Morgan et al., 1987). We felt it was appropriate to exclude it from the group mean data, as it likely did not represent a resting *c-fos* expression pattern. However, even if this dramatic outlier was included in the analysis, there still was no statistically significant difference between the two genotypes in the basal level of granule cell activation, as measured by *c-fos* expression (total c-fos+ cell density with outlier included: WT = 6210 ± 610 cells/mm^3^, KO = 68390 ± 60960 cells/mm^3^, n = 8,8 mice, p = 0.23). Thus, *CACNA2D2* KO mice do not appear to have grossly increased hippocampal activity when resting in their home cages.

### Altered hippocampal activity after induced seizures in *CACNA2D2* KO mice

*CACNA2D2* mutant mice can experience convulsive behavioral seizures after even gentle handling (Meier, 1968), and *CACNA2D2* KO mice have been observed to have convulsions in our animal facility with cage transfers. Thus, we considered the possibility that the KO mouse with widespread *c-fos* activation (Figure 1C) may have experienced a seizure involving the hippocampus due to handling shortly before euthanasia for tissue harvest. In the experiment above, mice were typically euthanized in fewer than 15 minutes after leaving their home cages in the animal facility. Thus, the majority of mice may not have had ample time to express detectable hippocampal *c-fos* after possible handling-induced convulsions, since *c-fos* takes approximately 60 minutes to achieve peak expression after activation (Morgan et al., 1987). To explicitly assess a potential relationship between handling and hippocampal activation, we euthanized WT and KO mice exactly one hour after gentle handling to determine whether handling impacts hippocampal granule cell activity.

Mice were picked up by the base of their tails, observed for 5 seconds, gently scruffed by the nuchal fur, and then placed in an observation cage, in a manner similar to our typical cage transfers prior to euthanasia, with the only difference being continuous observation for one hour prior to euthanasia. Although WT mice immediately resumed typical resting and exploratory behaviors after handling, *CACNA2D2* KO mice exhibited convulsive-like behavior similar to prior descriptions of handling-induced convulsions in spontaneous *CACNA2D2* mutant (*ducky*) mice (Meier, 1968; Snell, 1955), in which mice would at times rear and fall over, with bilateral limb clonus. In some mice, the convulsive behaviors would intermittently pause for minutes at a time, at which point the mice appeared to be resting quietly, after which similar convulsions would reappear. This behavior appeared distinct from the continuous stage 1/2-type seizures characteristic of pilocarpine-induced status epilepticus (Shibley & Smith, 2002) as there did not appear to be any convulsive behavior, or even tremor, during resting periods.

One hour after gentle handling, WT mice had c-fos+ expression patterns very similar to WT mice euthanized in our initial control experiments above (see Figures 1A,D; total c-fos+ cell density: WT immediate harvest = 6210 ± 610 cells/mm^3^, WT handled and delayed harvest = 5380 ± 1090 cells/mm^3^, n = 8,4 mice, p = 0.81). Surprisingly, however, despite clear convulsive activity (Figure 1E), *CACNA2D2* KO mice had dramatically reduced dentate *c-fos* expression one hour after handling when compared with WT littermates (total c-fos+ cell density: WT handled and delayed harvest = 5380 ± 1090 cells/mm^3^, KO handled and delayed harvest = 1540 ± 640 cells/mm^3^, n = 4,4 mice, p = 0.023, Figure 1D,F). KO mice that were handled also had significantly fewer c-fos+ cells than KO mice which were immediately harvested (n = 8,4; p < 0.005), suggesting that the decreased granule cell activity was a consequence of handling. Thus, not only was post-handling convulsive activity not associated with increased hippocampal *c-fos* expression, but our data demonstrated that handling actually dramatically decreased hippocampal activity. This suggests that the widespread *c-fos* activation in the single outlier animal above (Figure 1C) likely did not result from handling, and thus may have had a spontaneous seizure involving a separate mechanisms of neural circuit activation.

In addition to spontaneous and handling-induced behavioral seizures, *CACNA2D2* KO mice have a markedly decreased seizure threshold in response to low doses of the convulsant pentylenetetrazol (PTZ) (Ivanov et al., 2004). As behavioral seizures resulting from gentle handling were paradoxically associated with decreased activity in the dentate gyrus (Figure 1), we examined whether low-dose PTZ-induced seizures might also be associated with altered hippocampal activity in KO mice. All mice received a sub-threshold PTZ dose (30 mg/kg i.p.), which typically does not cause seizures in WT mice, but causes seizure-like activity in a majority of *CACNA2D2* KO mice (Ivanov et al., 2004).

As previously reported (Ivanov et al., 2004), while WT mice displayed little to no behavioral response to the subthreshold PTZ dose, *CACNA2D2* KO mice consistently responded to this dose with convulsive seizure activity (Figure 2A,B; WT: 0 of 9 mice with seizure score ≥ 3; KO: 8 of 8 mice with seizure score ≥ 3). Unlike the handling-induced convulsions, which were associated with decreased dentate *c-fos* activation, KO mice had significantly increased *c-fos* activation in the dentate gyrus after PTZ (Figures 2A,C). This cohort of mice expressed a somewhat bimodal distribution, in which half (4 of 8) of the KO mice had clearly detectable *c-fos* expression in nearly every granule cell, while the 3 of the remaining 4 mice had c-fos+ cell densities near, but still above, the mean for WT mice after PTZ. There was no clear association between the denser *c-fos* expression and post-PTZ seizure scores. In apparent contrast with the reduced *c-fos* expression after handling, these data suggest that the reduced behavioral seizure threshold in response to PTZ is accompanied by increased functional hippocampal network recruitment during these events.

**Figure 2.**
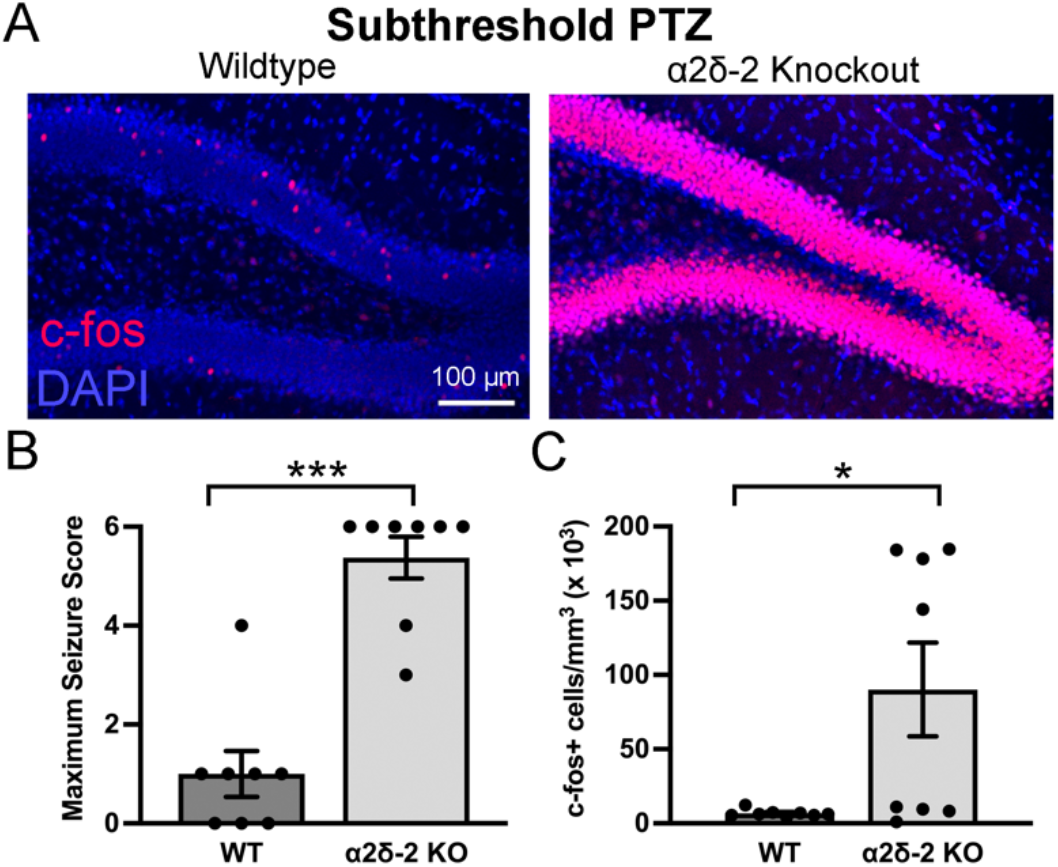
*CACNA2D2* KO mice have a reduced threshold for pentylenetetrazol-induced granule cell activation. A. Dentate *c-fos* expression (red) after low-dose (30 mg/kg) pentylenetetrazol (PTZ) administration in WT and KO mice. B. Maximum seizure intensity after PTZ (n = 8, 9; *** p < 0.0001). C. c-fos+ cell density after PTZ (n = 8, 9; * p = 0.028).

### Increased hilar mossy cell density without CA3 cell loss in *CACNA2D2* KO mice

Human TLE is characterized by numerous histopathological changes in the hippocampus, many of which are recapitulated in both chemoconvulsant and genetic mouse models of epilepsy (Loscher, 2017). Although handling-evoked convulsions can occur in *CACNA2D2* KO mice without clear hippocampal involvement (Figure 1D-E), hippocampal activation during seizures was clearly evidenced by the increased *c-fos* expression within the hippocampal granule cell layer after subthreshold PTZ administration (Figure 2) and was also observed on at least one occasion in an otherwise seemingly unprovoked manner (Figure 1C). Thus, we considered the possibility that functional and structural changes in the hippocampus might exist in these mice, and potentially contribute to the dentate hyperexcitability. We examined whether *CACNA2D2* KO mice demonstrated histopathological changes in the hippocampus similar to those frequently observed in mouse models of temporal lobe epilepsy (Sharma et al., 2007), beginning with the loss of excitatory cell populations in the dentate hilus and CA3 pyramidal cell layer.

The density of glutamatergic hilar mossy cells was quantified by staining for the calcium-buffering protein calretinin, a selective marker for excitatory mossy cells within the dentate hilus (Blasco-Ibanez & Freund, 1997). As calretinin (CR) is also expressed by immature granule cells approximately 1-2 weeks after mitosis (Brandt et al., 2003), we excluded small CR+ somata in the subgranular zone that were most likely immature granule cell neurons, and only counted large CR+ cells within the hilar region. Although hilar mossy cell loss is a prominent characteristic of many mouse models of epilepsy (Scharfman, 2016), we found that not only was the mossy cell population intact within the dentate hilus of *CACNA2D2* KO mice, but there was a small statistically significant increase in mossy cell density in KO mice (Figures 3A-B). Although this relatively small (38%) increase is of uncertain functional/biological significance, it is at least clear that mossy cell density was not reduced in KO mice at the age of spontaneous behavioral seizure onset.

**Figure 3.**
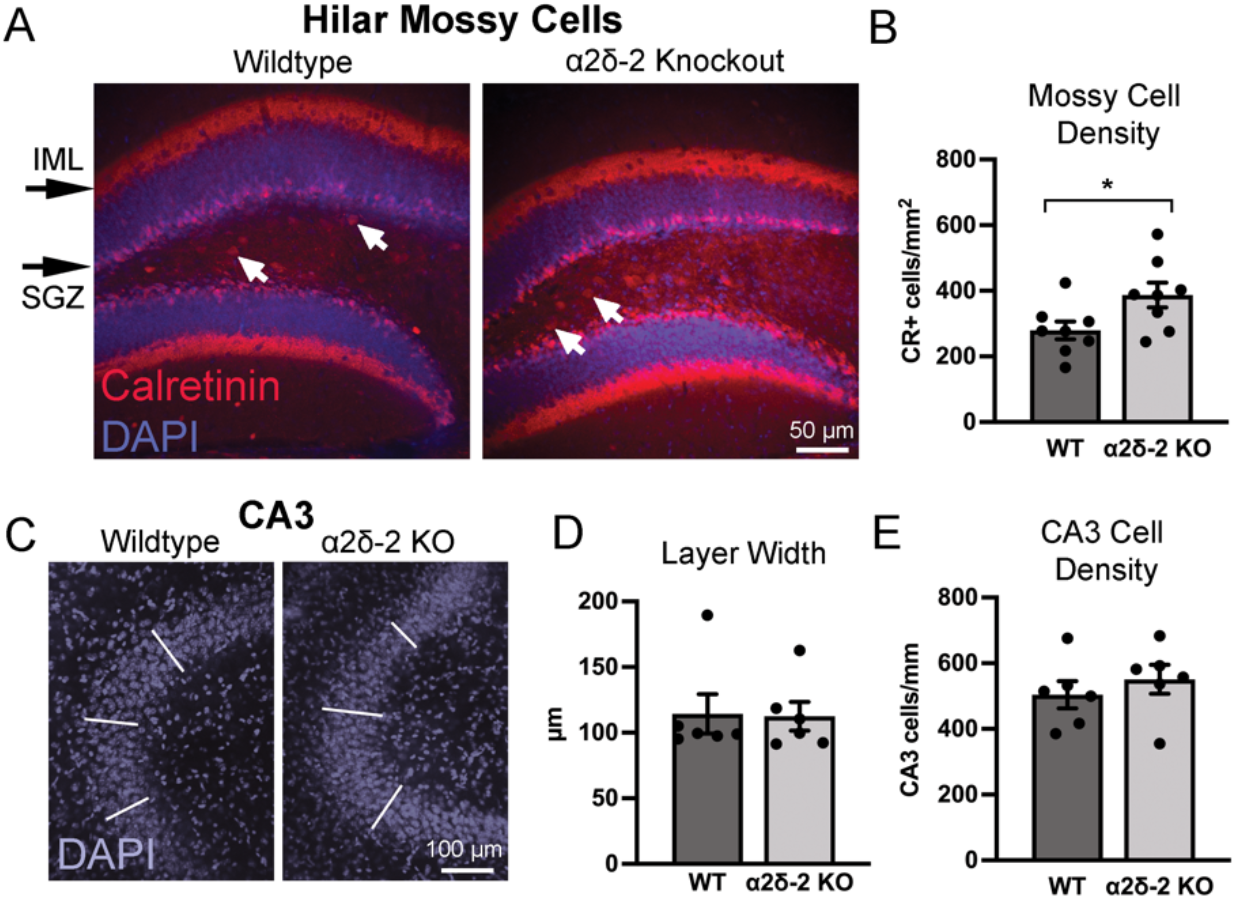
Hilar mossy cell and CA3 cell densities in *CACNA2D2* KO mice. A. Hilar mossy cells were identified as calretinin-positive cells (red; arrows) located within the hilar space. Mossy cell axons are visualized as a calretinin-stained band in the inner molecular layer (IML) outside of the granule cell layer (blue, DAPI). B. Quantification of hilar mossy cell density; small calretinin-positive immature granule cells in the subgranular zone (SGZ) were not included in the quantification. (n = 8, 8; * p = 0.037). C. CA3 pyramidal cell layer seen in cross section, as visualized using DAPI nuclear stain (blue). White lines designate 3 separate locations at which CA3 pyramidal layer width was measured. D. Quantification of CA3 pyramidal cell layer width (n = 6, 6; p = 0.94). E. Quantification of CA3 pyramidal cell layer density (n = 6, 6; p = 0.46).

Similarly, degeneration of hippocampal CA3 pyramidal cells occurs in numerous mouse models of epilepsy as well as in pathological specimens from human patients, which over time can contribute to an atrophied, sclerotic hippocampus (de Lanerolle et al., 2012). To assess CA3 cell loss, we quantified both CA3 pyramidal cell layer width and CA3 pyramidal cell density by quantifying DAPI-stained nuclei in hippocampal sections. Neither of these metrics was different between *CACNA2D2* WT and KO mice (Figures 3C-E), indicating that CA3 pyramidal cell loss is not a characteristic of *CACNA2D2* KO mice at this age.

### *CACNA2D2* KO mice do not exhibit retrograde mossy fiber axon sprouting

One pathognomonic histopathologic hallmark of temporal lobe epilepsy involves the retrograde sprouting of granule cell axons, also known as the mossy fibers (Nadler, 2003). Although the mechanisms driving mossy fiber sprouting are unclear, these fibers form recurrent excitatory circuits and may contribute to the initiation or propagation of seizures (Dudek & Shao, 2004; Hendricks et al., 2019; Tauck & Nadler, 1985).

To assess whether *CACNA2D2* KO mice manifest mossy fiber sprouting, we stained hippocampal sections for Zinc Transporter 3 (ZnT3) protein, which is highly expressed in mossy fiber terminals in both healthy and epileptic tissue (Murphy et al., 2011). In WT animals, ZnT3 was detected as dense staining throughout the dentate hilus and among the proximal dendrites of CA3 pyramidal cells (stratum lucidum), consistent with the known locations of mossy fiber synapses in healthy mice. KO mice had the same distribution of ZnT3-stained terminals (Figure 4), and most notably, a lack of ZnT3-stained puncta within the dentate inner molecular layer (IML), where spouted mossy fiber terminals are found in epileptic tissue (Murphy et al., 2011). No sections from any mouse had consistently observed ZnT3 positive boutons localized within the GCL or IML across sections (Figure 4), and a blinded quantification based on a modified Timm’s scoring system for mossy fiber sprouting showed no difference between groups (modified Timm’s score: WT = 0.52 ± 0.19, KO = 0.13 ±0.09, n = 6 per group, p > 0.05).

**Figure 4:**
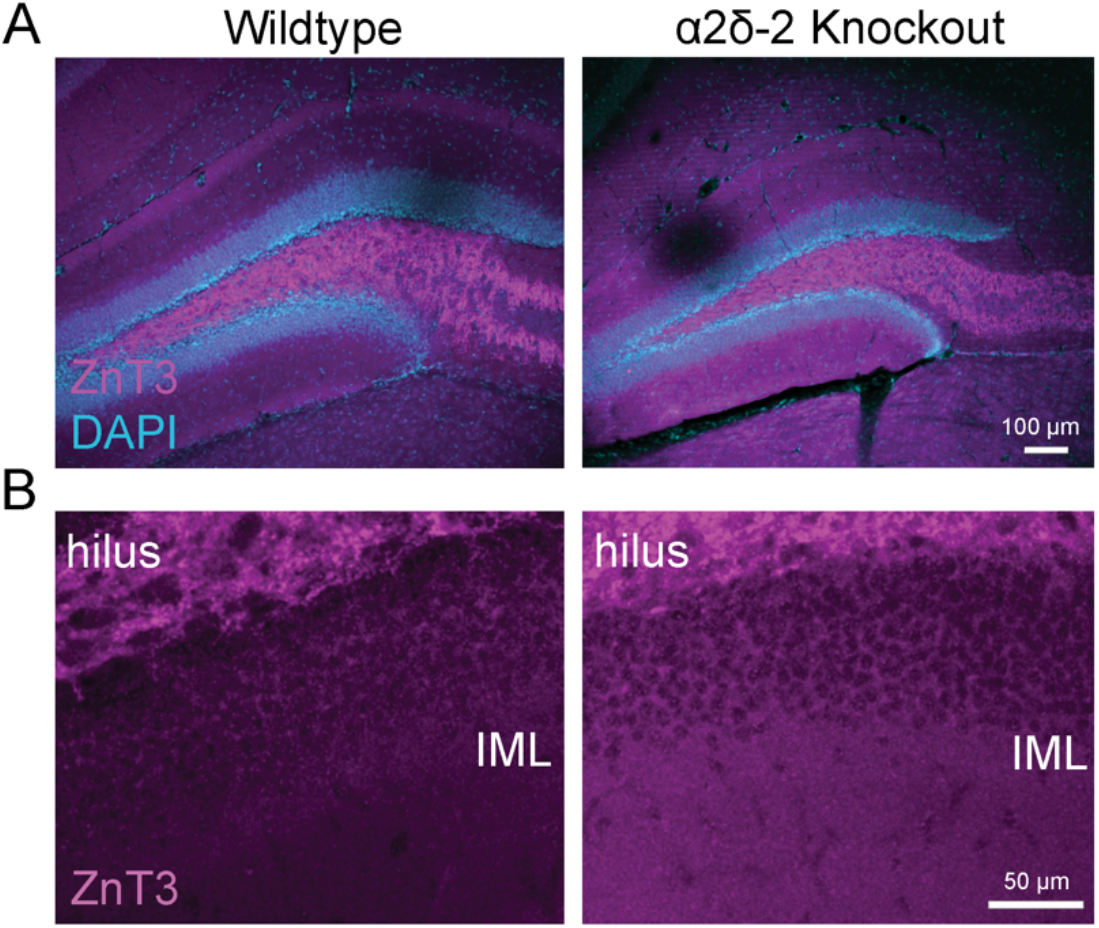
Mossy fiber sprouting is absent in *CACNA2D2* KO mice. A. Granule cell mossy fiber axon terminals in WT and KO mice were identified by staining for the mossy fiber terminal marker, ZnT3 (magenta). Granule cell layer highlighted with DAPI (blue). B. ZnT3-stained terminals were exclusively found within the hilar space, and not within the inner molecular layer, which would have indicated pathologic retrograde sprouting.

### Granule cell proliferation and hippocampal neurogenesis are not increased in juvenile *CACNA2D2* mutants

The hippocampal dentate gyrus is characterized by ongoing granule cell neurogenesis in adulthood, and the rate of hippocampal adult neurogenesis is modulated by numerous biological contingencies, including neurological disease (Eisch et al., 2008; Gage, 2002). Seizures dramatically affect hippocampal neurogenesis in the dentate, causing increased neural stem cell proliferation, as well as accelerated dendritic outgrowth, outward cell body migration, and increased synaptic innervation of newly generated neurons (Overstreet-Wadiche et al., 2006; Parent et al., 1997). The strong association of seizures with aberrant neurogenesis has led to the suggestion that aberrant neurogenesis may directly contribute to the pathogenesis of epilepsy (Cho et al., 2015; Danzer, 2012; Parent & Kron, 2012).

To quantitatively assess hippocampal neurogenesis in juvenile *CACNA2D2* KO mice, we first stained hippocampal sections for the nuclear marker, Ki67. Ki67 is expressed by cells undergoing active mitosis (Scholzen & Gerdes, 2000) and commonly used as a measure of cell proliferation. In hippocampal sections from both WT and KO mice, Ki67+ nuclei were found within/near the dentate sub-granular zone (SGZ), as would be expected for proliferating neural stem cells from which granule cells are generated, which reside in this region (Figure 5A). There was no difference in the density of Ki67+ labeled cells between groups (Figure 5C), indicating that *CACNA2D2* KO does not alter cell proliferation within the neurogenic niche of the dentate gyrus.

**Figure 5.**
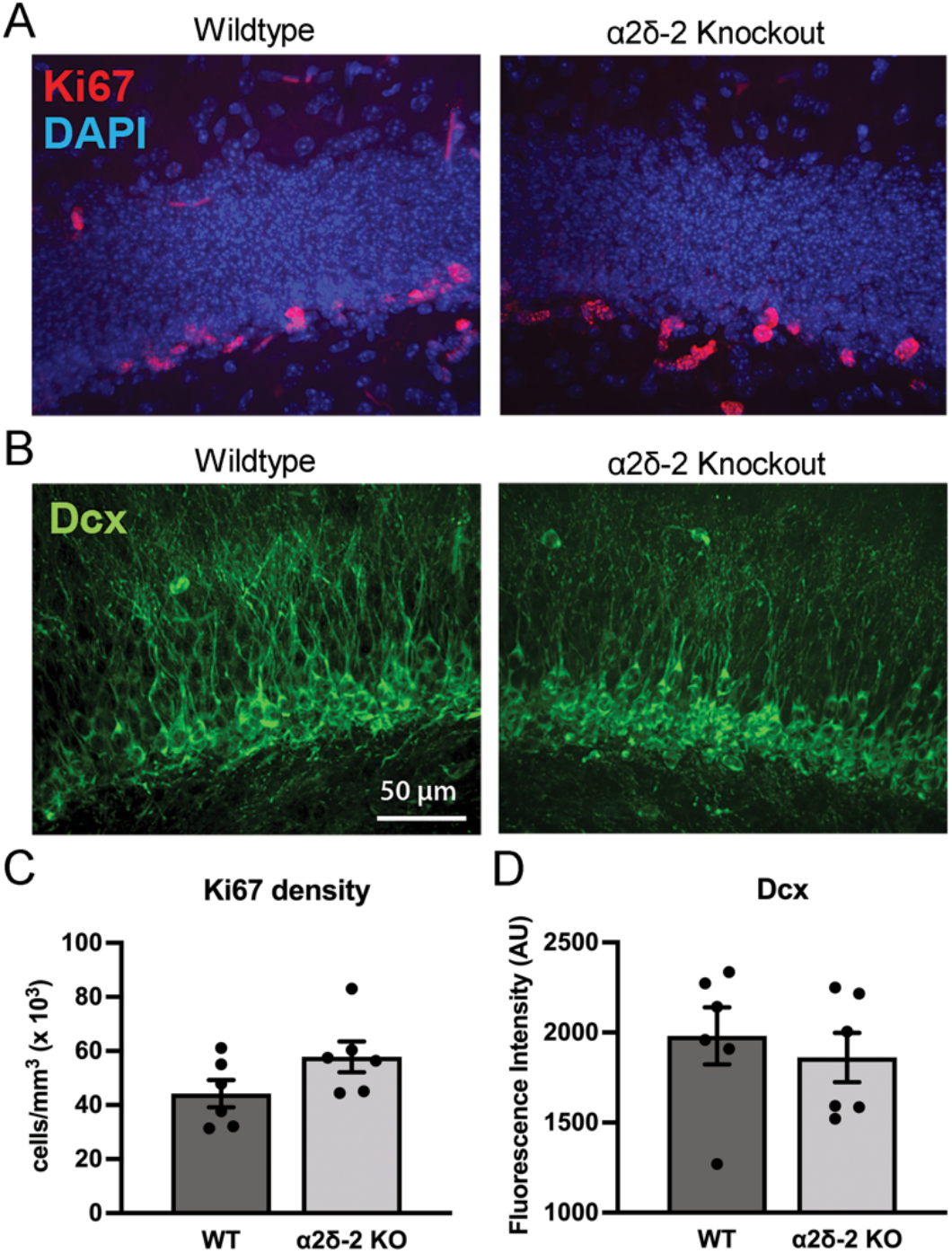
Hippocampal cell proliferation and granule cell neurogenesis are unchanged in juvenile *CACNA2D2* KO mice. A. Mitotic cells in the dentate subgranular zone identified by the proliferation-associated nuclear marker Ki67 (red) in WT and KO mice; granule cell layer nuclei were counterstained with DAPI (blue). B. Immature granule cell neurons identified by expression of the microtubule-associated protein doublecortin (Dcx, green) in WT and KO dentate sections. C. Quantification of Ki67+ cell density (n = 6, 6; p = 0.11). D. Quantification of Dcx+ cell density using staining fluorescence intensity in the granule cell layer (n = 6, 6; p = 0.58).

As Ki67 is not selective for neurogenic precursor cells, we stained hippocampal sections for doublecortin (Dcx), a microtubule-associated protein that is expressed by immature granule cell neurons early after mitosis. Again, both genotypes had similarly dense appearing Dcx+ staining (Figure 5B), indicative of the robust neurogenesis that is typical of juvenile mice at this age. Quantification of Dcx fluorescence demonstrated no significant difference between groups (Figure 5D), suggesting no difference in the density of immature, recently born granule cells in KO mice. Together with a lack of difference in cell proliferation, these data indicate that *CACNA2D2* KO mice do not have gross abnormalities in dentate granule cell neurogenesis at this age.

### Parvalbumin-positive interneuron density is unchanged in *CACNA2D2* KO mice

α2δ-2 is highly expressed in Parvalbumin-positive (PV+) interneurons (Cole et al., 2005), which mediate feed-forward inhibition and control neuronal excitability in the hippocampus. In some models of epilepsy, PV+ interneuron density in the hippocampus is reduced, leading to the suggestion that deficits in inhibitory circuits might contribute to neuronal circuit hyperexcitability (Dudek & Sutula, 2007). Thus, to assess whether loss of α2δ-2 protein is associated with a loss of PV+ interneurons, we stained hippocampal sections from WT and *CACNA2D2* KO mice for PV.

In the dentate gyrus, PV+ cells were sparse and predominantly localized along the SGZ, the typical location of PV+ basket-type interneurons, as well as more rarely within the hilus and molecular layer. To focus on the basket-type PV cells most closely associated with feed-forward inhibition in the dentate gyrus, we separately quantified PV+ cells within the granule cell layer and dentate hilus. The PV+ cell density in both regions was the same between WT and KO mice (Figures 6A-C).

**Figure 6.**
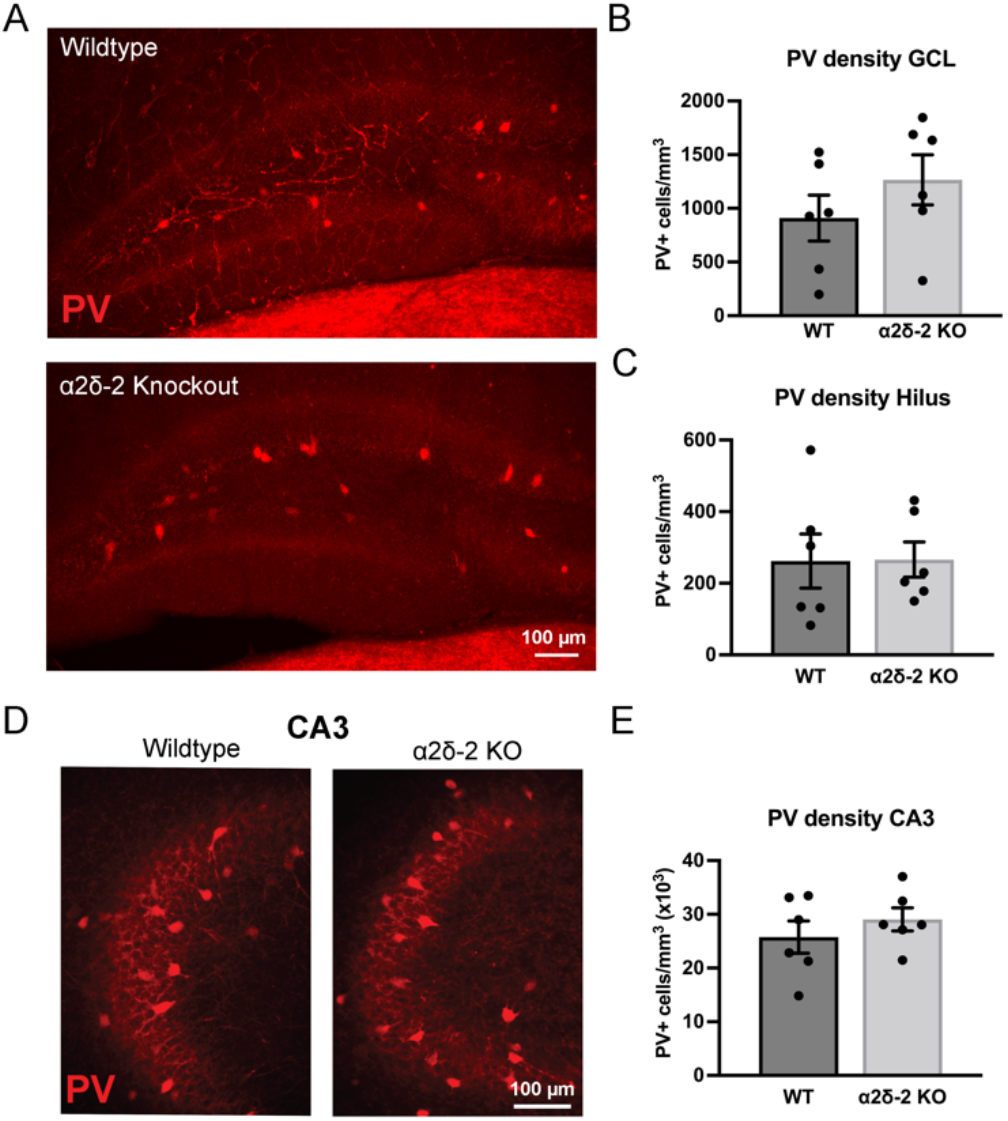
Parvalbumin interneuron staining in WT and *CACNA2D2* KO mice. A. Confocal stack image projections of parvalbumin-stained (PV, red) dentate gyrus tissue from WT and KO mice. B, C. Quantification of PV+ cell density within both the granule cell layer (GCL; B; n = 6, 6; p = 0.29) and dentate hilus (C; n = 6, 6; p = 0.97). D. Confocal stack image projections of parvalbumin-stained (PV, red) hippocampal CA3 pyramidal cell layers from WT and KO mice. E. Quantification of PV+ cell density in the CA3 pyramidal cell layer (n = 6, 6; p = 0.40).

As a change in PV+ cell density in CA3 could profoundly alter hippocampal circuit excitability, we also quantified PV+ cell density within the CA3 pyramidal cell layer. Again, there was no difference in the PV+ cell density within the CA3 pyramidal cell layer between WT and KO mice (Figures 6D-E). Additionally, subjective assessments of cell morphology performed on WT and KO images after unblinding did not reveal any obvious difference in cell structure or axonal projections of PV+ neurons. Together, these data indicate that *CACNA2D2* KO is not associated with gross structural changes within the hippocampal PV+ interneuron cell network.

### Glial marker staining is not increased in *CACNA2D2* KO mice

Epilepsy is associated with glial activation (Dossi et al., 2018). Thus, neuronal hyperexcitability can occur as a direct consequence of neuronal injury, or alternatively may be due to underlying increases in neuroinflammation that could secondarily contribute to epileptogenesis (Wetherington et al., 2008). Both astrocytes and microglia modulate neural circuit function and structure, through both direct interactions with neurons and synapses and indirectly through the release of neuromodulatory and neurotropic factors (Christopherson et al., 2005; Ekdahl et al., 2009; Schafer et al., 2013). Thus, we immunohistochemically evaluated two hippocampal glial cell populations, astrocytes and microglia, in brain tissue from WT and *CACNA2D2* KO mice.

To determine whether hippocampal astrocytes are altered in number or activation state in the absence of α2δ-2 protein, we stained hippocampal sections for Glial Fibrillary Acidic Protein (GFAP), which both identifies astrocytes and increases in expression with astrocytic activation (Schiffer et al., 1986). GFAP+ cells with typical astrocytic spider-like morphology were found throughout the dentate molecular, granule cell, and hilar regions in both WT and KO mice, and also included GFAP+ radial processes within the granule cell layer, which resembled previously described radial glial stem cells. Quantification of the density of GFAP+ cells between WT and KO mice revealed no difference in the density of astrocytes between WT and KO mice (Figures 7A-B). Additionally, as GFAP expression can increase in activated astrocytes, we compared immunohistochemical signal intensity between groups, and also found no differences between groups (WT = 359 ± 14 A.U.; 367 ±16 A.U.; n = 6,6; p = 0.70).

**Figure 7.**
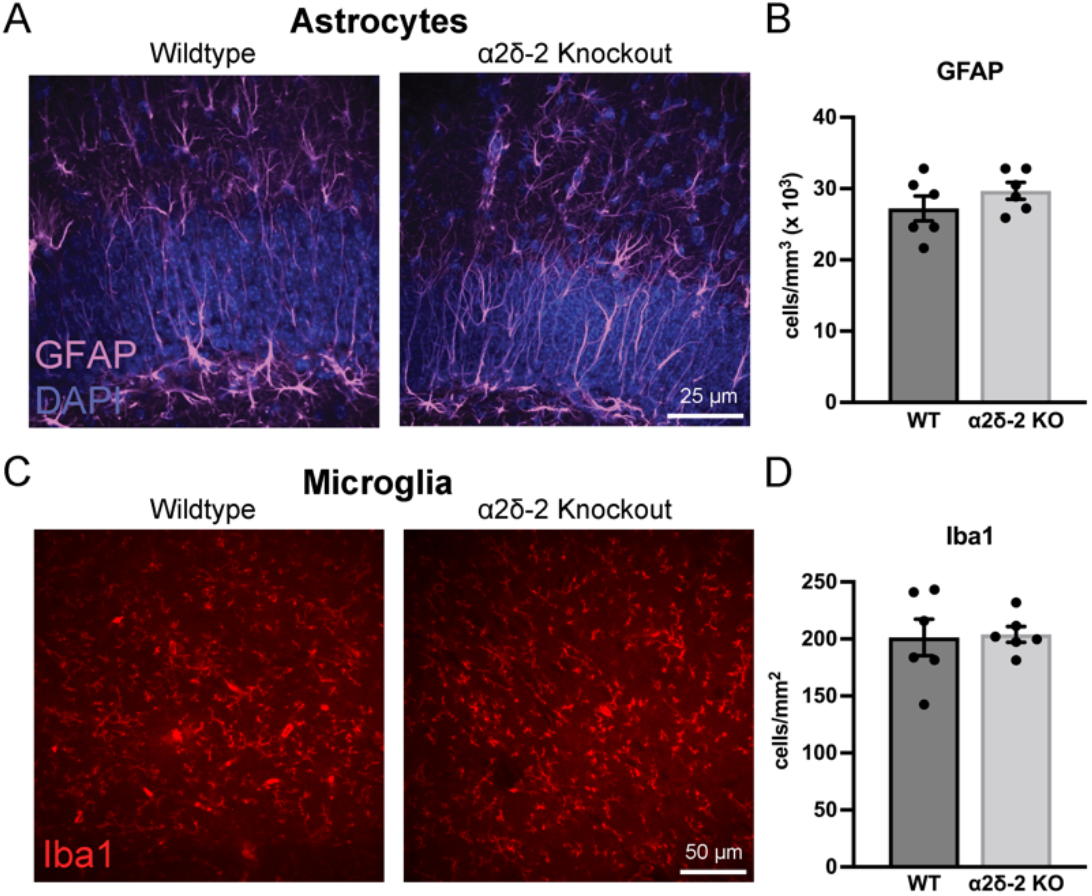
Glial activation is not changed *CACNA2D2* KO mice. A. Confocal stack image projections of glial fibrillary acidic protein (GFAP)-stained dentate granule cell layers (DAPI, blue) from WT and KO mice. B. Quantification of GFAP+ cell density across both the entire image field (n = 6, 6; p = 0.29). C. Confocal image of Iba1-stained (red) dentate molecular layer tissue from WT and KO mice. D. Quantification of Iba1+ cell densities (n = 6, 6; p = 0.88).

As microglia powerfully modulate neuronal circuits and play a role in secondary neuronal damage, we stained hippocampal tissue for the microglial marker Iba1. There was no difference in the density or distribution of microglia (Iba1) between WT and KO mice (Figures 7C-D). Similarly, there was no difference in the density of activated microglia, defined as those staining for the marker Mac-2 (Galectin-3), which was similarly low in both WT and KO mice (Mac-2+ cells in WT = 698 ± 206 cells/mm^3^; KO = 1221 ± 238 cells/mm^3^; n = 6,6 mice, p = 0.13). Together with the lack of change in GFAP immunohistochemistry, these data indicate that loss of α2δ-2 protein is not associated with pathophysiological hippocampal glial cell proliferation or activation.

## Discussion

Mice lacking α2δ-2 protein have increased behavioral seizure susceptibility as well as seizure-associated differences in hippocampal granule cell activity. Surprisingly, hippocampal recruitment during seizure episodes in *CACNA2D2* KO mice differed remarkably after two different seizure-inducing stimuli, gentle handling and subthreshold pentylenetetrazol injections, which either dramatically decreased or increased hippocampal granule cell recruitment respectively. Presumably these differences relate to different mechanisms underlying the behavioral responses to two different seizure-provoking stimuli, and likely involve distinct patterns of neural circuitry activation in association with seizure events.

### Reduced threshold for hippocampal activation in the absence of α2δ-2 protein

*CACNA2D2* KO mice manifest tonic-clonic seizures in response to doses of PTZ that are subthreshold for WT mice (Ivanov et al., 2004), and we now confirm that a subthreshold dose of PTZ also substantially increases hippocampal activity in KO mice when compared with WT mice. The reduced ability to maintain network homeostasis when challenged with proconvulsant stimuli, combined with no change in baseline levels of *c-fos* activation, suggests potential dysfunction in inhibitory feedforward networks that maintain network quiescence during periods of enhanced activation in KO mice. Since one of the major interneuron subtypes mediating this form of inhibition, PV+ interneurons, is not diminished in density, a plausible explanation for this phenotype could involve PV+ neuron dysfunction (without a clearly associated gross structural phenotype) or alterations in the excitatory drive of PV+ neurons.

α2δ-2 was initially identified as a VGCC subunit, suggesting that α2δ-2-expressing cells, such as hippocampal PV+ cells (Cole et al., 2005), might manifest functional changes in both Ca^++^ signaling and Ca^++^-dependent subcellular processes that are not reflected in structural analyses. Indeed, cerebellar Purkinje cells, which highly express α2δ-2 in WT mice, demonstrate profound functional differences in both their afferent synaptic input as well as Ca^++^-dependent action potential afterhyperpolarization in the absence of α2δ-2 protein (Beeson et al., 2020; Beeson et al., 2022). Similarly, neurons lacking the α2δ-1 isoform have altered neurotransmitter release phenotypes (Hoppa et al., 2012), which might be recapitulated in cells that rely on α2δ-2 for similar functions. Either cell intrinsic (action potential waveform) or synaptic (GABA release or excitatory innervation) dysfunction in PV+ cells could cause seizures without grossly apparent structural phenotypes, and might also preferentially manifest during conditions of enhanced circuit activation (e.g. PTZ). Additionally, these phenotypes could also become more prominent as circuits mature during mouse adolescence/weaning, when afferent connectivity into the hippocampus has reached adult levels. As genetic epilepsy syndromes can involve functional (electrophysiological) deficits without clear structural correlates (Meisler et al., 2001), hippocampal function in *CACNA2D2* KO mice could be significantly compromised without structural changes in a similar manner.

### Paradoxically reduced granule cell activation during handling-induced convulsions in *CACNA2D2* KO mice

Surprisingly, *CACNA2D2* KO mice had reduced *c-fos* expression in dentate granule cells in association with handling-induced convulsions, indicating that activity in the hippocampus had actually decreased below basal levels during these seizure events. This not only suggests that this form of seizure activity in KO mice is mediated by brain regions outside of the hippocampus, but also that compensatory mechanisms may actually produce a form of peri/post-ictal depression of activity in the hippocampus, which might be similar to prior descriptions of post-seizure reductions in cellular activation in the brain (Morgan et al., 1987). The anatomical location involved in generating and propagating handling-associated seizure activity in these mice is not known, but prior observations provide some potential insights.

Early histopathological studies of α2δ-2 mutant (*ducky*) mice identified cerebellar, hindbrain, and spinal cord abnormalities, including Purkinje cell atrophy, reduced volumes and cell sizes in brainstem nuclei, and axonal lesions in several spinal cord motor pathways (Meier, 1968). A future comprehensive analysis of post-handling *c-fos* expression could elucidate whether any of these regions might have been involved in post-handling seizures. A more recent analysis identified reduced neocortical size and reduced cortical layer V thickness with increased cell density in *ducky* mice (Geisler et al., 2021). As this cortical layer contains pyramidal cells that both receive inputs from and project to the thalamus, it may well be the anatomical substrate for the electroencephalographic spike-wave discharges (SWDs) characteristic of homozygous mutant mice (Barclay et al., 2001). However, as these SWDs are characteristic of “absence” type seizures typically associated with behavioral arrest, rather than motor convulsions (Barclay et al., 2001), it remains unclear whether the observed motor convulsions in *CACNA2D2* KO mice are the result of aberrant thalamocortical activity or due to circuit dysfunction elsewhere in the brain or spinal cord.

One particular difference that was uncovered in our study was a small but statistically significant increase in the density of calretinin-positive hilar mossy cells. Mossy cells preferentially drive inhibitory circuit function and can control seizure activity (Bui et al., 2018). Thus, an increased density of functional mossy cell outputs could potentially reduce granule cell activity levels in KO mice, if convulsive seizures originate outside of the hippocampus and increase inhibitory circuit activity through an enhanced mossy cell network. However, given the small absolute magnitude of the increased hilar mossy cell density, it is unclear whether this small increase would substantially alter functional output of this pathway, as mossy cells project widely to bilateral hippocampus and might homeostatically adjust their output connectivity accordingly (Butler et al., 2022).

### Future Directions

We performed our analyses at one month of age, which correlates with the onset of behavioral seizures noted in mice lacking α2δ-2 protein (Ivanov et al., 2004; Meier, 1968). The lack of several histopathological markers of temporal lobe epilepsy at this time indicates that the circuit rearrangements that they represent (recurrent mossy fiber sprouting, loss of hippocampal neuron populations, glial activation) are not required for mice to express seizures or increased excitability at this age.

Although behavioral seizures do not require global dentate gyrus activation in the hippocampus at this early timepoint, seizures can lead to secondary changes in circuit structure and progressively worsening disease (Cavazos et al., 1991). Thus, our analysis does not preclude potential hippocampal involvement in the progression and worsening of epilepsy as mice continue to age, which might include changes in hippocampal structure at adult mice. Improvements in animal husbandry that allow for enhanced survival of KO mice into adulthood (Geisler et al., 2021) could facilitate analysis at these later stages.

An inherent limitation to our study is the restriction of our analysis to a limited number of histologic assays. For example, while our focus on PV+ interneurons was derived in part from the expression of α2δ-2 protein in this cell type, other interneuron types, such as somatostatin-expressing interneurons, also decrease in density in certain forms of epilepsy (Hofmann et al., 2016; Santhakumar et al., 2000) and were not examined here. Finally, electrophysiologic studies will hopefully elucidate any changes to intrinsic hippocampal circuit function, as well as any potential alterations in afferent connectivity into the hippocampus.

Future work will hopefully determine whether restoration of α2δ-2 function later in life, perhaps through viral vector-mediated rescue approaches, would be sufficient to diminish behavioral seizures in the face of seizure-related circuit remodeling, which could be relevant to families with α2δ-2 mutations as well as others with epilepsy-associated channelopathies.

## Acknowledgments

We thank Gary Westbrook and his lab members for helpful discussions of our data, Stefanie Kaech and the OHSU Advanced Light Microscopy Core for imaging assistance, Drs. Sergey Ivanov and Lino Tessarollo for generously providing mutant mice, and members of the Schnell lab for feedback and support.

